# Transcriptomic analysis of organotypic porcine retina cultures

**DOI:** 10.64898/2026.04.16.718959

**Authors:** Siavash Khosravi, Grazia Giorgio, Federica Staurenghi, Tanja Schoenberger, Peter Gross, Margit Ried, Julia Frankenhauser, Sebastian Eder, Elke Markert, Remko A. Bakker, Sepideh Babaei, Nina Zippel

## Abstract

Porcine organotypic retinal explant cultures are widely used to study retinal neurodegeneration under controlled conditions, but the biological process that occurs in the retinal explant over time due to preparation-induced injury and culture are not well understood. Here, we generated a time-resolved transcriptomic reference for porcine neural retinal explants-maintained ex vivo for 10 days. Global expression profiles are strongly separated by culture time, with Day 0 clearly distinct from cultured samples and at Day 7 and Day 10 showing the highest similarity, indicating a transition toward a later stabilized state. Across the time course, 3,187 genes were differentially expressed relative to Day 0, with the largest shifts occurring at an early stage of culture (Day 1–Day 3). Pathway-level analyses revealed coordinated remodeling involving inflammatory signaling, and metabolic/bioenergetic changes, including reduced mitochondrial and oxidative phosphorylation–related programs at later time points. Here, we provide a time-resolved transcriptomics reference dataset for cultured porcine retinal explants. These data can build a foundation to interpret data generated in this model, differentiate changes inherent to the explant culture from treatment-specific effects and to select appropriate experimental windows for mechanistic studies of retinal degeneration.

## 1. Introduction

Retinal degenerative and vascular diseases remain leading causes of irreversible visual impairment worldwide and progress in therapeutic development depends strongly on experimental models that capture the cellular diversity, layered architecture, and metabolic demands of the mature human retina^1,2^. Ex-vivo retinal organotypic (explant) cultures can bridge the gap between simplified cell culture systems and in vivo animal models by largely preserving key aspects of native retinal tissue organization, cellular architecture and intercellular interactions, making them highly relevant for studying retinal biology under controlled experimental conditions while reducing the need for animal use. At the same time, organotypic cultures inherently impose defined stresses (e.g., axotomy, altered oxygen/nutrient diffusion, loss of systemic cues), that drive time-dependent molecular and cellular changes characteristic for neurodegeneration. Consequently, retinal explants are widely used as model systems for retinal degeneration, particularly for investigating mechanisms of inflammation, gliosis, apoptosis, neurodegeneration and related therapeutic interventions. To correctly interpret effects of experimental interventions within the context of the model’s intrinsic degenerative processes, it is essential to have a clear temporal understanding of how culture-induced molecular programs dynamically evolve over time^2^.

Among available large-animal systems, the pig (Sus scrofa) has emerged as a highly relevant translational model in retinal research because many aspects of ocular anatomy and retinal morphology resemble humans more closely than commonly used rodent models^1 3^. Pigs have a large eye size, a cone-enriched central retina organized as a visual streak that is used as a macula-analog in translational studies ^4^. Photoreceptor mapping studies further demonstrate a structured spatial organization and cone distribution in pig retina that supports its use as a surrogate system for human retinal research^5^. Consistent with this, pigs are increasingly used for retinal disease modeling and preclinical evaluation of therapeutic concepts, including interventions where retinal size and architecture strongly influence feasibility and outcome measures^6^.

Ex-vivo porcine retinal explants enable medium-throughput testing in a controlled environment while maintaining layered tissue structure^7^. Recent methodological work has established that porcine retinal explants can be maintained over extended periods (up to several weeks) with measurable preservation of retinal cell markers and morphology, and that culture conditions and handling strongly influence photoreceptor preservation^7 8^. Despite these advances, a major limitation remains: the time-dependent remodeling that occurs during porcine retinal explant culture is not yet comprehensively understood, making it difficult to define an optimal experimental window or to separate culture-induced programs from treatment-specific effects ^17^.

In this study, we performed bulk RNA sequencing on porcine neural retinal explants cultured ex-vivo for up to 10 days, sampling tissue at Day 0, Day 1, Day 3, Day 7, and Day 10. Our primary aim was to map the transcriptomic trajectories that unfold during explant maintenance in culture, thereby providing a molecular framework for interpreting porcine retinal explants as an experimental model system. Importantly, despite the wide-spread use of porcine explants in translational retinal research, a comprehensive, time-resolved reference of culture-driven transcriptomic remodeling is not available. By generating and organizing these data across multiple culture time points, we tried to elucidate the transcriptional changes that assist investigators to identify appropriate experimental time windows. In parallel, we aimed to define the early versus late molecular changes associated with culture adaptation and stress, including immune-like signaling, glial and inflammatory response programs, alterations in neuronal/synaptic processes, and shifts in metabolic and bioenergetic pathways. Together, this analysis provides a structured transcriptomic reference that strengthens the interpretability, comparability, and translational utility of the porcine explant model for studies of retinal degeneration.

## 2. Results

To assess the similarity of gene expression profiles between samples and replicates, Pearson correlation coefficients were computed across all samples and visualized as a correlation heatmap with unsupervised hierarchical clustering (Figure 1A). Replicates within each time point showed high within-group correlation and samples segregated into major time-associated blocks. Samples of Day 0 formed a distinct cluster with high internal correlation compared to other samples (Figure 1A). Samples of Day 1 and Day 3 formed separate clusters as well. Importantly, samples of Day 7 and Day 10 showed higher mutual similarity, appearing as a combined late-time block relative to the earlier time points (Figure 1A). These patterns indicate that global transcriptomic similarity is highest among replicates of the same time point and among late time points (Day 7/Day 10) compared with earlier transitions.

**Figure 1.**
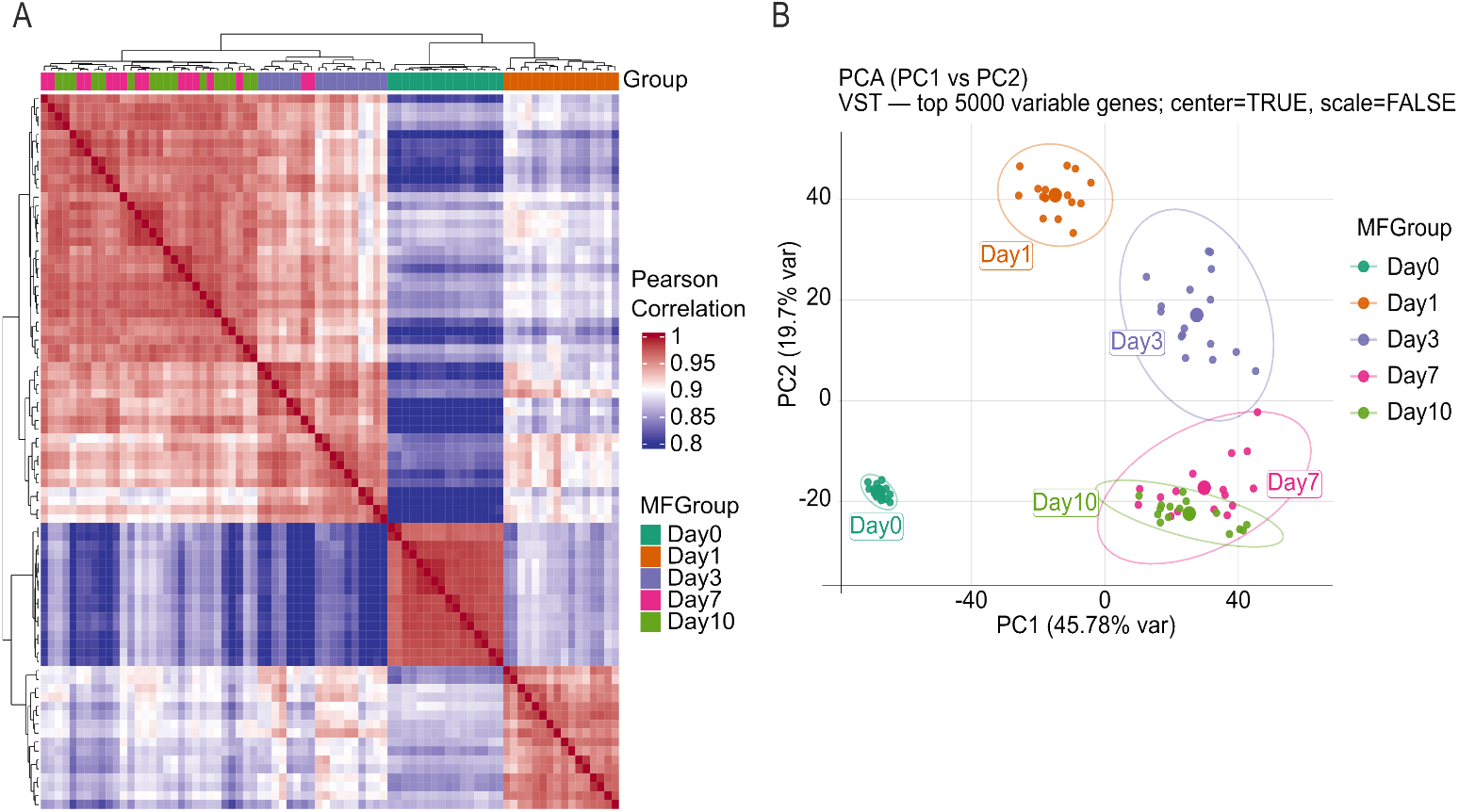
Global transcriptomic similarity across the porcine retinal explant culture time course. (A) Sample–sample Pearson correlation heatmap calculated on variance-stabilized (VST) expression values using the top 5000 most variable genes. Samples are annotated by culture time point (MFGroup: Day 0, Day 1, Day3, Day 7, Day 10) and ordered by unsupervised hierarchical clustering (complete linkage; correlation-based distance). (B) Principal component analysis (PCA) performed on the same VST-transformed expression matrix (top 5000 variable genes; center = TRUE, scale = FALSE). Each point represents one sample colored by time point; axes indicate the percentage of variance explained (PC1: 45.78%, PC2: 19.7%).

To further evaluate sample similarity, principal component analysis (PCA) was performed on variance-stabilized (VST) expression values using the top 5,000 most variable genes to assess global sample relationships across the culture time course (Figure 1B). Samples clustered strongly by time point, indicating that incubation time was the dominant factor determinant for transcriptomic variation in this dataset. These results are In agreement with the patterns which were observed in Pearson correlation analysis (Figure 1A). The first two principal components explained 45.7% (PC1) and 19.7% (PC2) of the variance, respectively (Figure1B). The PCA plot clearly showed a direct temporal ordering. For instance, Day 0 samples formed a tight cluster distinct from all cultured samples, while Day 1 and Day 3 each formed separate clusters distanced from Day 0 (Figure1B). Day 7 and Day 10 samples clustered closest to one another and showed overlap, indicating higher similarity between late time points compared with earlier stages (Figure 1B). Similar time-dependent divergence of mouse retinal explants during ex-vivo culture has been reported in organotypic retina systems, where culture duration is a major determinant of tissue state and readouts^9^.

To understand transcriptional changes driving the molecular and physiological changes during the culture time, differential gene expression (DGE) analysis was performed. DGE analysis revealed a total of 3,187 unique genes that were differentially expressed across the culture time course (Log2FC > 1, adjusted P value < 0.05). When classified by baseline contrasts (Day 1/Day 3/Day 7/Day 10 vs Day 0), as it is depicted in figure 2A, the number of differentially expressed genes increased in Day 1 and Day 3. However, the number of differentially expressed genes decreased on Day 7 and Day 10, most notably due to a reduction in the number of downregulated transcripts in Day 7 and Day 10 (Figure 2A). In line with the PCA analysis (supplementary Figure 1), the number of differentially expressed genes between Day 3 and Day 1, Day 7 and Day 3 and, Day10 and Day 7, were reduced drastically. This might indicate a drastic transcriptional shift at beginning of the culture that tapers off over time.

**Figure 2.**
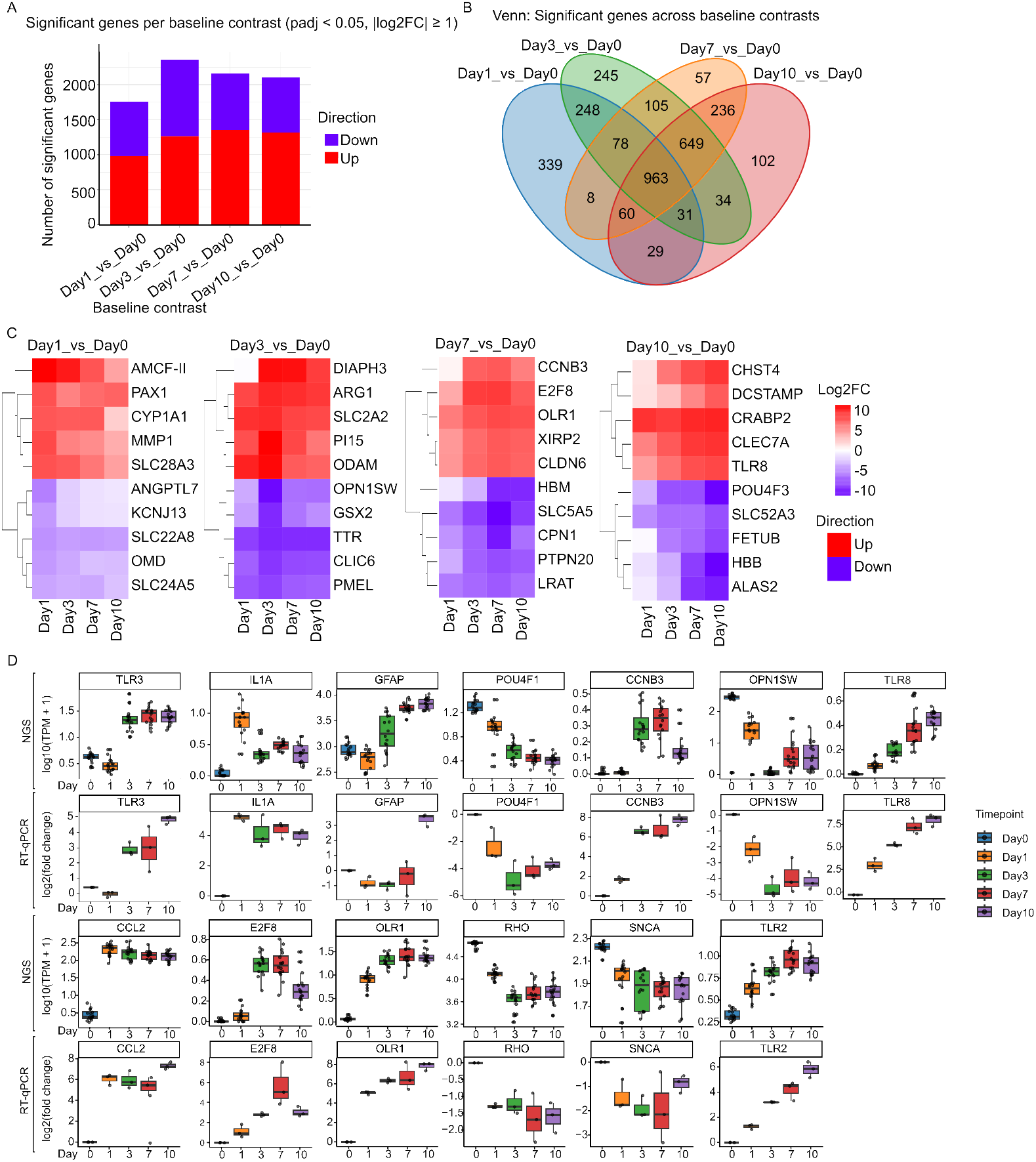
Differential gene expression dynamics in porcine retinal explants across culture time (baseline contrasts). (A) Venn diagram showing overlap of significantly differentially expressed genes identified in each baseline comparison (Day 1/Day 3/Day 7/Day 10 vs Day 0). (B) Number of significant DGEs per baseline contrast, separated by direction (Up/Down). Differential expression was assessed using DESeq2, and significance was defined as Benjamini–Hochberg adjusted p-value (FDR) < 0.05 with an effect-size threshold of |log2FC| ≥ 1. (C) Top regulated genes (five most upregulated and five most downregulated) for each baseline contrast, displayed by log2 fold change (log2FC). (D) Quantification of mRNA levels. For each indicated gene, mRNA levels were measured by RT-qPCR and compared to RNA levels obtained from NGS analysis across all time points. The upper panel displays RNA levels from the NGS analysis in log_10_(TPM + 1). The lower panel presents mRNA quantification using TaqMan probes for each time point in log_2_(fold change).

The Venn diagram (Figure 2B) shows that each contrast contains both time-point–specific DGE and a shared component overlapping across contrasts, consistent with the time-dependent sample segregation observed in Figure 1 (PCA and Pearson correlation clustering). In total, 963 DGEs were shared among all four baseline contrasts (Figure 1B).

We computed differential gene expressions (DGE) for each baseline comparison (Day 1/Day 3/Day 7/Day 10 vs Day 0) and, within each contrast, selected the genes showing the largest LogFC changes (Figure 2C). AMCF-II, PAX1, CYP1A1, MMP1, and SLC28A3 were among the most upregulated genes in Day 1. While we identified downregulated genes including ANGPTL7, KCNJ13, SLC22A8, OMD, and SLC24A5. In Day 3 vs Day 0 comparison, DIAPH3, ARG1, SLC2A2, PI15, and ODAM were upregulated, while OPN1SW, GSX2, TTR, CLIC6, and PMEL were downregulated. For Day 7 vs Day 0, CCNB3, E2F8, OLR1, XIRP2, and CLDN6 were upregulated, whereas HBM, SLC5A5, CPN1, PTPN20, and LRAT were downregulated. In the Day 10 vs Day 0 contrast, CHST4, DCSTAMP, CRABP2, CLEC7A, and TLR8 were identified as upregulated, while POU4F3, SLC52A3, FETUB, HBB, and ALAS2 were among the most significant downregulated genes.

Additionally, we measured mRNA levels of selected genes by RT-qPCR in an independent sample set and made a comparison with our NGS data to assure validity of our NGS analysis (Figure 2D). Thirteen mRNAs level were measured and compared with their corresponding RNA level in our NGS analysis (Figure 2D). Among thirteen mRNA levels which were quantified, all of them demonstrate identical trend in comparison to the RNA level in our NGS analysis. Lastly, we measured mRNA levels of same genes in the early hours after the dissection. Notably, we observed that mRNA level of some genes alters way earlier than the others (supplementary Figure 2) like ILA1.

To deepen our understanding of longitudinal pathway activity, we quantified enrichment of Gene Ontology Biological Process (GO-BP) gene sets using GSVA scores among samples. We tested time-dependent differences with a limma linear modeling framework which is applied directly to the GSVA score matrix. This workflow tests differential pathway activity and controlled multiple testing using FDR adjustment. Figure 3A demonstrates the top five significantly upregulated and top five significantly down-regulated GO terms for each time-point contrast (FDR < 0.05).

**Figure 3:**
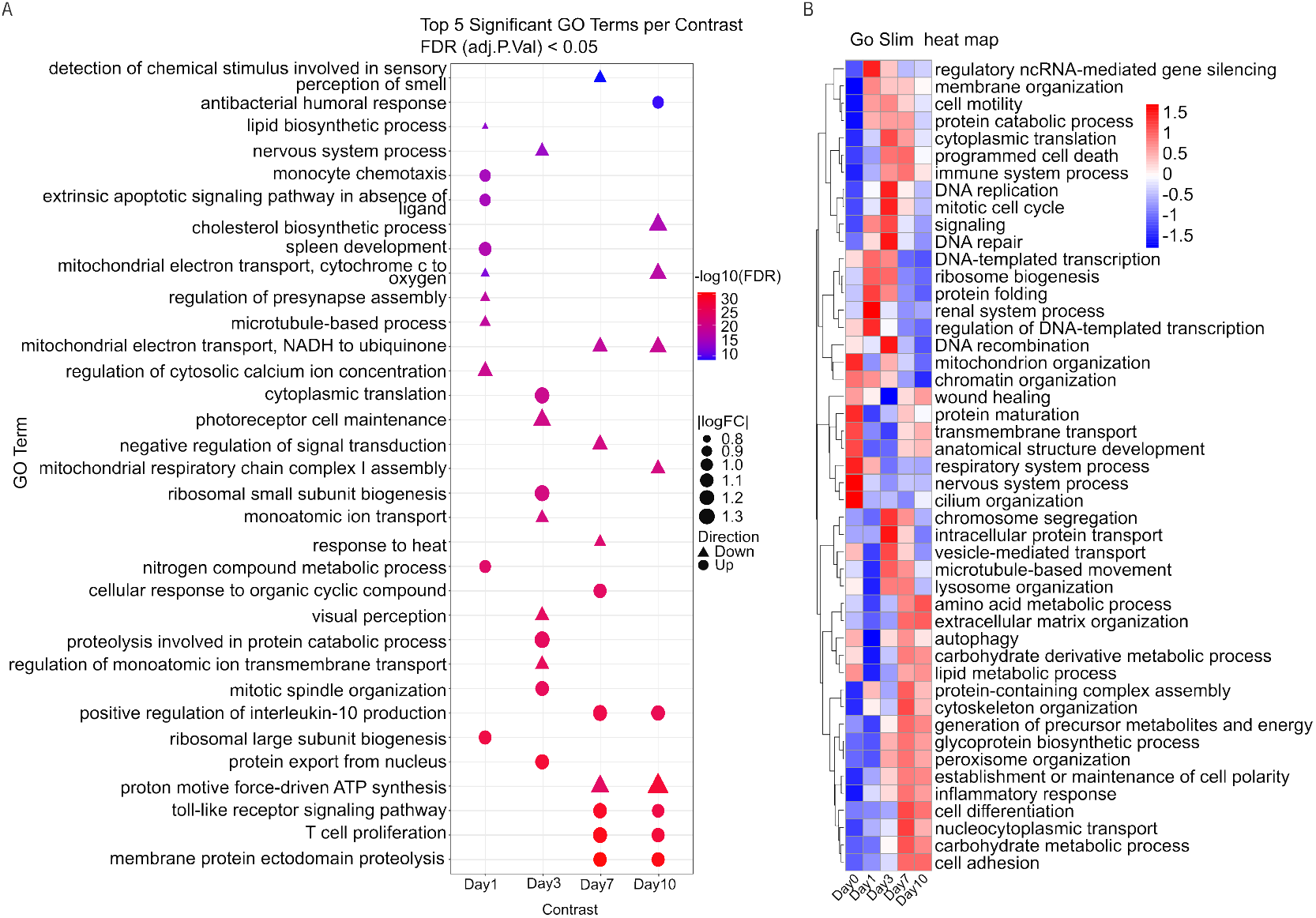
Pathway-level remodeling during porcine retinal explant culture. (A) Dot plot summarizing the top five significantly upregulated and top five significantly downregulated GO Biological Process gene sets for each baseline contrast (Day 1/Day 3/Day 7/Day 10 vs Day 0). Dot color reflects the strength of significance (−log10(FDR)). the direction of change (up/down), and dot size indicates the LogFC. Triangle represents downregulation of a pathway, and circle represents upregulation of a pathway. Pathway differences were tested on GSVA scores using a linear-modeling framework with Benjamini–Hochberg FDR < 0.05. (B) Heatmap of significantly altered GO Slim biological process categories across Day 0–Day 10, derived by mapping significant GO terms to higher-level GO Slim classes and visualizing row-scaled (z-scored) mean GSVA activity per time point.

Among contrast, the most prominent GO-BP signals fell into two biologically axes: immune activation/regulation and core cellular metabolism/homeostasis. In Day 7 and 10 Immune-associated terms are among the strongest pathway-level shifts (Figure 3A). This includes toll-like receptor signaling pathway, T cell proliferation, monocyte chemotaxis, antibacterial humoral response and positive regulation of interleukin-10 production (Figure 3A).

Additionally, several top-ranked GO terms reflected changes in bioenergesis and biosynthesis. These included mitochondrial pathways (mitochondrial respiratory chain complex I assembly, mitochondrial electron transport, NADH to ubiquinone, mitochondrial electron transport, cytochrome c to oxygen, proton motive force-driven ATP synthesis), translation-related programs (cytoplasmic translation, ribosomal small subunit biogenesis, ribosomal large subunit biogenesis) and proteostasis involved in protein catabolic process, membrane protein ectodomain proteolysis). Lipid-related terms (cholesterol biosynthetic process and lipid biosynthetic process) and apoptosis-related signaling (extrinsic apoptotic signaling pathway in absence of ligand) also appeared among the top differential GO terms,

Notably, we identified GO terms linked to ion transport and stimulus/sensory-associated biology, including regulation of monoatomic ion transmembrane transport, monoatomic ion transport, regulation of cytosolic calcium ion concentration, visual perception, photoreceptor cell maintenance, regulation of presynapse assembly, nervous system process and detection of chemical stimulus involved in sensory perception of smell. It is worth noticing that GSVA identified the cholesterol biosynthetic process as a strongly downregulated GO term at Day 10. This is in accordance with our differential gene expression results, where OLR1 (oxidized low-density lipoprotein receptor 1) was among the most strongly upregulated genes at Day 7.

To have higher-level view of pathway dynamics that complement the more detailed GO-term analysis, we performed the GO Slim. All GO terms were subsequently mapped to higher-level GO Slim categories using the GO-Slim generic subset and GO hierarchical relationships. The result is shown in the heatmap (Figure 3B) summarizing biological processes that change significantly over time (FDR ≤ 0.05)^3,4^. Additionally, the heatmap reveals a clear stage-specific remodeling rather than a uniform response across all days. Notably, several broad GO Slim categories overlap with our previous analysis of topmost significant enriched GO terms. This overlap includes immune activation, translation/proteostasis, mitochondrial/energy metabolism, and programmed cell death. A closer look at the heatmap reveals that they change in a coordinated manner. For instance, many signals are relatively low at early stages (Day 0/Day 1), become strongly elevated around Day 3, and remain partially elevated or are reorganized by Day7–Day10.

In consistency with our previous results, immune axis is a significant broad GO term (Figure 3B). Immune system processes and inflammatory responses increase toward Day 1 and Day 3, while the inflammatory signature remains elevated through Day 7–Day 10. These findings are in line with the enrichment of top significant GO terms of toll-like receptor signaling pathway, T cell proliferation, monocyte chemotaxis, and antibacterial humoral response (Figure 3A).

Additionally, GO-Slim analysis identified the metabolic and cell-fate themes as another significant axis in line with our previous results. Translation/proteostasis categories such as cytoplasmic translation and protein catabolic process increase strongly around Day 3 and remain elevated by Day 7, consistent with the previous result of signals for ribosomal biogenesis/translation and proteolysis-related processes. In parallel, mitochondrial and energy-related categories display a time-dependent restructuring too. For instance, mitochondrion organization varies across Days, whereas generation of precursor metabolites and energy becomes more prominent at later stages particularly on Day 7. Finally, programmed cell death increases at later stages (notably Day 3–Day 7), mirroring the enrichment of apoptosis-related signaling and supporting the result of in significant GO term in GSVA dot plot.

Ultimately, we performed Gene set enrichment analysis (GSEA) on the DESeq2 Wald-statistic– ranked transcriptome across the culture time course. GSEA analysis identified a strong positive enrichment of the inflammatory response program (FDR = 2×10?5) for Day 1 (Figure 4A), indicating a coordinated shift toward innate immune activation with increasing time in explant culture. The leading-edge genes visible in the plot include canonical TLR adaptors and sensors (TIRAP, TICAM1, MYD88; TLR2n-induced injury and culture are not well understood. Here, we generated a time-resolved transcriptomi–MyD88/TRIF signaling driving NF-κB–linked cytokine/chem-okine induction in retinal cells and retinal disease contexts.

**Figure 4:**
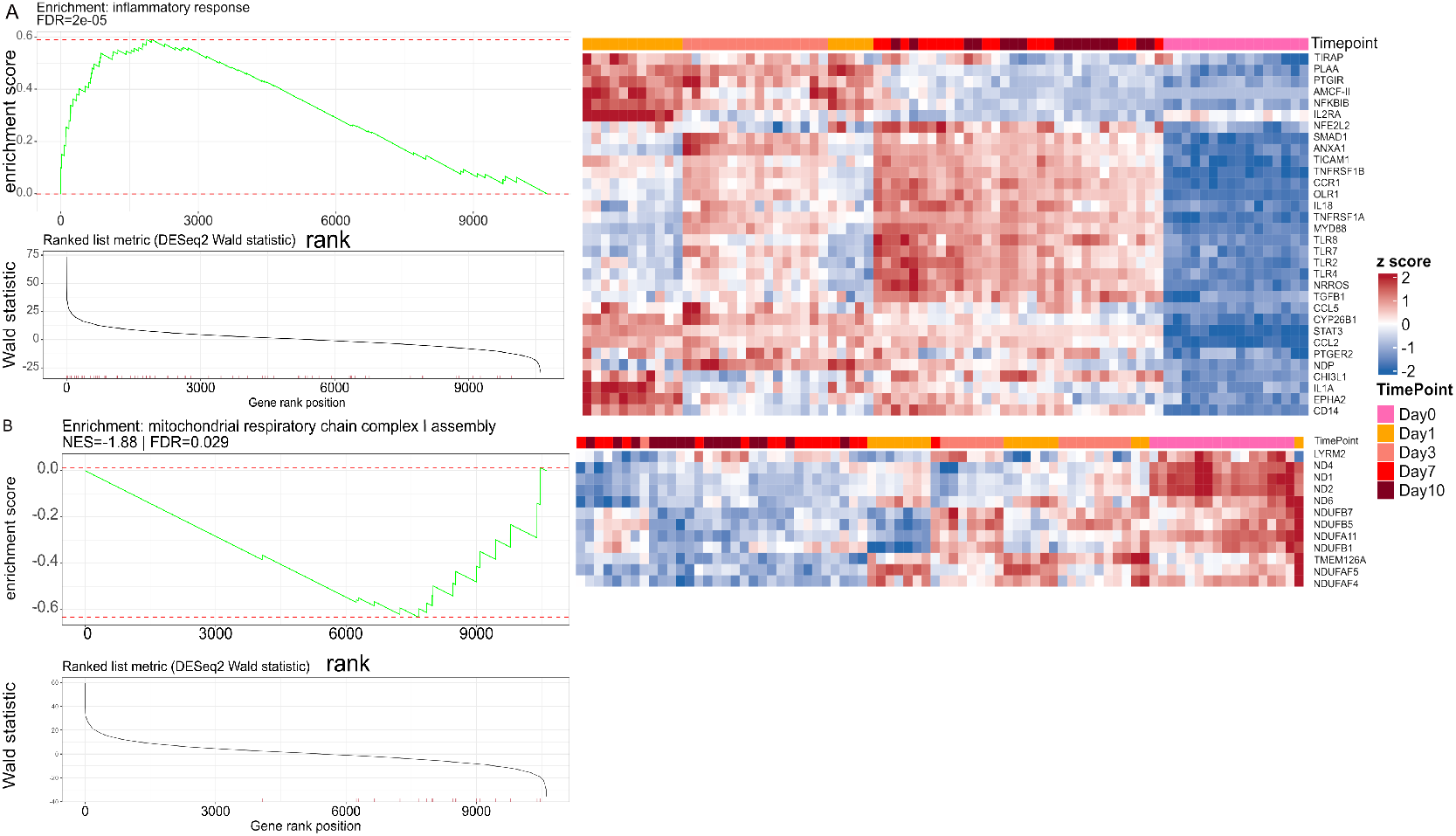
Gene set enrichment analysis (GSEA) identifies inflammatory activation and mitochon-drial remodeling during retinal explant culture. (A) GSEA for the inflammatory response gene set using a genome-wide ranking of genes by the DESeq2 Wald statistic for the baseline contrast Day 1 vs Day 0. The panel shows the enrichment plot (running enrichment score across the ranked list), the corresponding ranked-list metric, and a heatmap of genes contributing to the enrichment signal (leading-edge genes) across samples/time points. Significance is reported as Benjaminiß–Hochberg FDR. (B) GSEA for the mitochondrial respiratory chain complex I assembly gene set performed analogously for the baseline contrast Day 10 vs Day 0, displaying the enrichment plot, ranked-list metric (DESeq2 Wald statistic), and a leading-edge gene heatmap across samples/time points. Enrichment is summarized by normalized enrichment score (NES) and FDR.

In contrast, the mitochondrial respiratory chain complex I assembly gene set showed significant negative enrichment (NES = −1.88, FDR = 0.029) for Day 10 (Figure 4B), driven by reduced representation of core complex I components and assembly factors (e.g., ND1/ND2/ND4/ND6, NDUFB1/5/7, NDUFA11, TMEM126A, NDUFAF4/5). Because complex I is central to oxidative phosphorylation and is a major contributor to mitochondrial ROS when dysregulated, this pattern is consistent with progressive bioenergetic remodeling/attenuation during culture—mechanistically relevant to retinal neurodegeneration, where complex I deficiency can precede or coincide with innate immune/inflammatory transcript induction and neuronal vulnerability.

Our observation of immune activation revealed by GSVA and GSEA led us to study changes in the microglial population in retinal explants by immunostaining. We assessed two markers, Iba1 and Ki67, to study proliferating and non-proliferating myeloid cells (Figure 5A). Ki67 staining indicates active cellular proliferation, whereas Iba1 staining identifies cells of the microglial and macrophage lineages. We then quantified co-staining of Iba1 and Ki67. Cells co-stained for Iba1 and Ki67 indicate proliferating myeloid cells in the tissue, cells stained for Ki67 but not Iba1 imply proliferation of other retinal cell types and cells stained for Iba1 but not Ki67 represent the non-proliferating myeloid cell population. Our quantification clearly demonstrates that the numbers of proliferating cells increase starting from Day 7 and that part of the proliferating cell population are myeloid cells (Figure 5B). This finding is consistent with our previous analysis showing that cellcycle genes, including CCNB3 and E2F8, reach their peak at Day 10 (Figure 2B). More importantly, we also observed activation of innate immune receptors and myeloid-associated genes such as TLR8 and CLEC7A, indicating a prominent inflammatory state (Figure 2B, D). Additionally, we performed GFAP staining on samples across all time points, a marker expressed in astrocytes and activated Muller cells. Analysis of GFAP staining showed that gliosis increased and became widespread throughout the different layers of retinal tissue during the culture period, indicating specifically an activation of Muller cells that are spanning all retinal layers.

**Figure 5:**
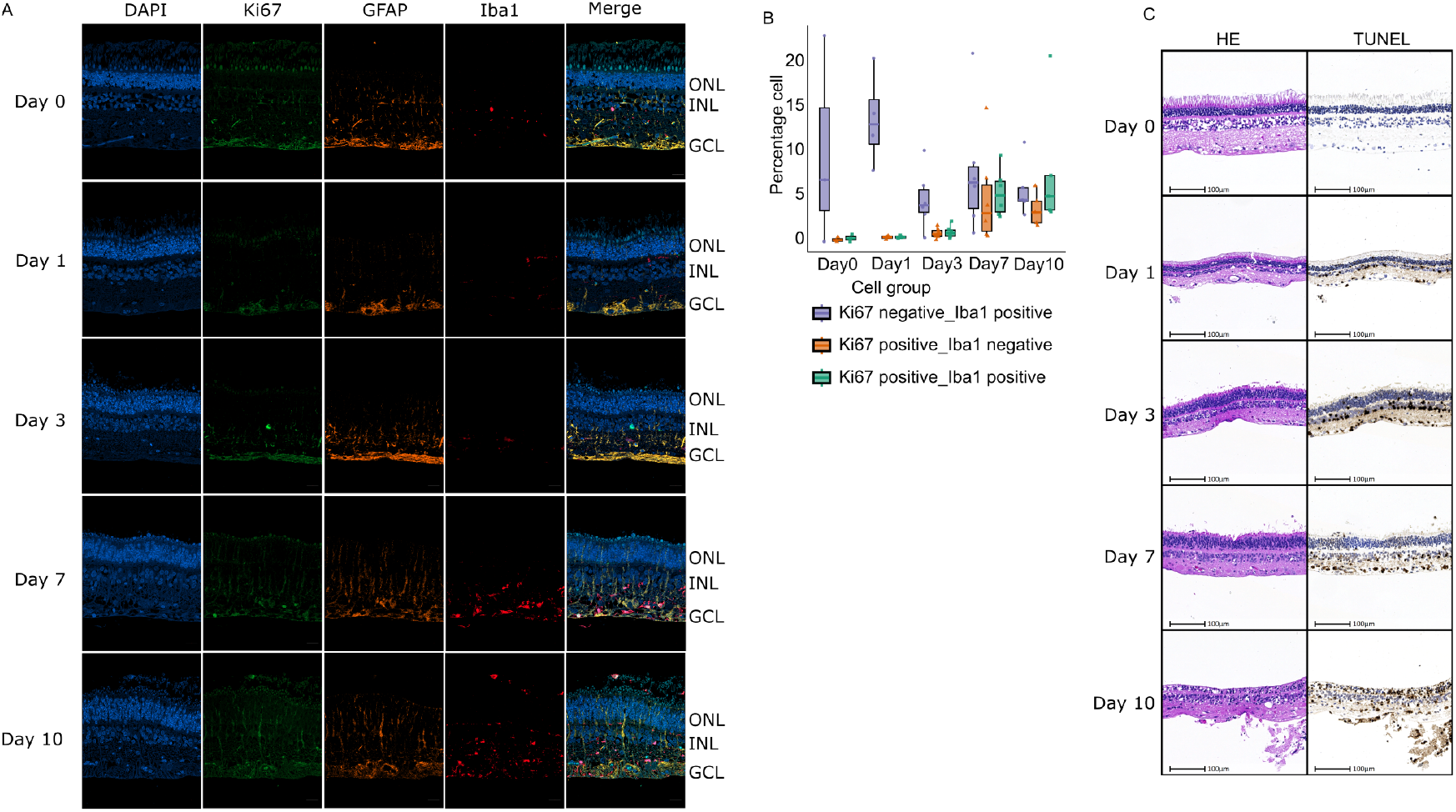
Immunofluorescence labeling and immunohistochemistry of retinal segments across different time points. (A) Representative fields show sections of mounted retinal explants at the indicated time points, co-stained for Iba1 (red), Ki67 (green), and GFAP (orange). (B) Quantification of the percentage of cells expressing different markers in panel A. Three different cell groups were measured, as indicated. (C) TUNEL assay in porcine retinal explants over the culture period. Serial sections were processed for H&E and TUNEL staining (brown TUNEL signal; nuclei counterstained with DAPI in blue).

To characterize degenerative changes and cell death in retinal explants during the culture period, Morphology was evaluated on hematoxylin and eosin (H&E) combined with TUNEL stained serial sections (Figure 5C).

On Day 0, no morphological changes were present. Slight vacuolization or disruptions, including (partial) absence of outer and inner segments of the photoreceptor layer were considered processing- and handling related supported by negative TUNEL stain (Figure 5C). From Day 1 of cultivation onwards, cell density in the ganglion cell layer is decreased in some samples and more than 50% of the ganglion cells as well as a subset of inner nuclear layer cells stain TUNEL positive (Figure 5C). Importantly, we have observed reduction of RBPMS staining in flat mounts of porcine retina at days 5 and day7 (supplementary Figure 3). This agrees with our observation of a sharp increase in the z-score of the GO-Slim term ‘extrinsic apoptotic signaling pathway in absence ligand’ at Day 1 (Figure 2B). While the number of TUNEL positive cells and vacuolization increased over culture time in all retinal layers, the cell density decreased, and thinning of plexiform layers occurred. Additionally, in samples with preserved inner and outer segments flattening and loss of photoreceptors can be observed. Besides, we observed a similar increase in the z-score of the GO-Slim term ‘programmed cell death’ On Day 7. Finally, we observed shrunken hyperchromatic nuclei and loss of retinal architecture in most samples on Day 10.

Consistent with our earlier differential expression analyses, the GO Slim time-course in Figure 6 describes a coordinated injury-response cascade in retinal explants following enucleation (Figure 6). Neuronal processes are sharply downregulated at Day 1 and remain suppressed through Day 10, indicating rapid loss of neuronal gene programs (Figure5). In parallel, immune pathways rise early peaking between days 3–7. This reflects an acute response that gradually resolves by Day 10. Programmed cell death shows a transient rise during the acute phase, preceding a later increase in cell-differentiation terms that peak around Day 7, consistent with tissue remodeling and reparative transcriptional states. Together, these dynamics integrate our earlier observations of an immediate neuronal shutdown followed by transient immune/inflammatory activation and subsequent engagement of differentiation programs.

**Figure 6:**
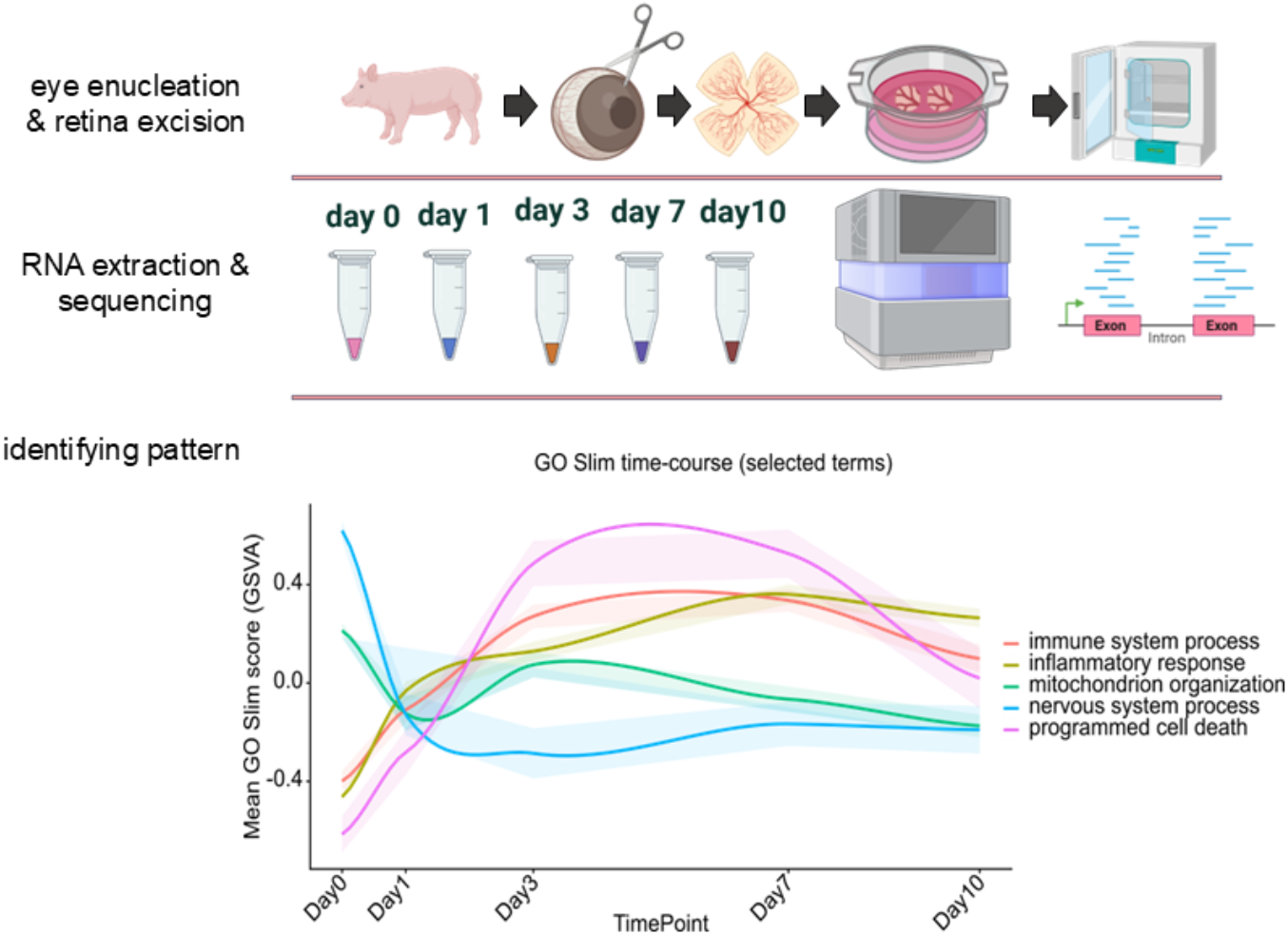
Schematic overview of the experimental and analytical workflow used to map culture-driven transcriptomic remodeling in porcine retinal explants. Enucleated porcine eyes were dissected to isolate neural retinal explants, which were maintained as organotypic cultures and sampled longitudinally (Day 0, 1, 3, 7, and 10). Total RNA was extracted from explants at each time point and subjected to bulk mRNA sequencing, followed by computational analysis to identify time-dependent expression patterns and pathway dynamics across the culture period. The figure summarizes the three major steps of the study: eye enucleation/retina excision, RNA extraction and sequencing, and pattern discovery from the transcriptomic time course (Biorender is used to prepare this figure).

## 3. Discussion

A central outcome of this study is that culture duration represents the primary factor that determines global gene expression dynamics in porcine retinal explants. This conclusion is supported by the strong within–time point concordance that is observed in Pearson correlation clustering, the clear temporal separation revealed by PCA and distinct time dependent biological processes by GSVA, GSEA and GO term analysis. Such behavior is fully consistent with the concept that organotypic retinal culture triggers an injury-and-adaptation program initiated at the time of enucleation and tissue preparation.

During explant preparation, axotomy inherently occurs, as removal of the retina from the eyeball damages retinal ganglion cell (RGC) axons at the optic nerve head. This injury initiates a cascade of cellular, molecular and functional changes across multiple retinal cell types. In addition, in retinal explant cultures the retina is separated from the retinal pigment epithelium (RPE), causing photoreceptors to lose physical contact with this supporting layer. Although pieces of RPE are cultured alongside to provide necessary growth factors, the metabolic and structural support normally provided by the RPE is disrupted and functions like i.e. photoreceptor outer segment phagocytosis and visual pigment recycling are lost. This ultimately compromises photoreceptor integrity, function and survival ^10^. Moreover, isolation of the retina also disconnects the retinal vasculature from the systemic circulation, abolishing perfusion and therefore oxygen and nutrient delivery and systemic signaling cues. Collectively, this factor, RGC axotomy, loss of RPE-photoreceptor interaction and disconnection of the vasculature, drive a progressive reshaping of the retinal transcriptome during the culture period. Comparable time-dependent progress has been reported for adult mammalian retinal organotypic cultures, where structural preservation may remain preserved while molecular and functional signatures diverge substantially with days of in-vitro culture^9^,^11, 12^.

Substantial shared DGE fraction across all baseline contrasts (Figure 2B) (including a core set present at every cultured time point) supports the existence of a conserved program due to damages during enucleation, retina excision and culture conditions. Differential expression analysis identified thousands of genes altered relative to Day 0, with the largest transcriptomic shift occurring in early days like Day 1 and Day 3. This is perfectly in line with our observation in PCA in which Day 1 and Day3 have a prominent clear separation of PCA1 and PCA2 with Day 0 compared to Day 7 and Day 10. This observation is also supported by Pearson correlation analysis in which Day 1, and Day 3 have a clear distinct block from Day 0 and Day 7/Day 10. A possible interpretation is that the transition from Day 0 to Day 1/3 represent the acute response to mechanical injury, axotomy, hypoxia/reoxygenation-like stress, and extracellular matrix disruption. Additionally, this agrees with comparative work demonstrating that retinal organotypic cultures share overlapping signatures of glial activation and neuronal stress across culture time, though the timing and magnitude of these responses can differ^9^,^11, 12^.

Tian et al. (2022) identified approximately 1,115 and 3,403 differentially expressed genes (DEGs) in mouse retinal ganglion cells (RGCs) on Days 1 and 3 after a crush injury ^13^. In line with this, our bulk analysis of porcine retinal explants revealed that Day 3 saw the most significant transcriptomic changes, with the majority of the 2,354 DEGs emerging early. This consistency across species and experimental models indicates that enucleation-induced RGC axotomy triggers a shared acute injury response.

In later days of culture, close clustering of Day 7 and Day 10 samples suggests that following an early phase of rapid transcriptomic remodeling (Day 0–Day 3) the explant transcriptome enters a more stable state characterized by slower incremental changes. This is supported by the marked reduction in the number of DGEs between adjacent late time points (Day 7 vs Day 10) demonstrating a strong overlap of Day 7/10 in our PCA analysis suggesting that late-stage transcriptomics are more similar to each other than to early-stage cultures. This indicates that by Day 7–Day 10, the explants enter a later culture state rather than an extension of the early response. At this stage, the tissue may have partly adjusted to ex-vivo conditions and shifted toward a glia-dominated and inflammatory profile (e.g., gliosis and microglial activation), together with metabolic reprogramming as we observed in our GSVA and GO-Slim analysis ^9, 11^,^12^.

Investigation of the most strongly regulated genes by time point (Figure 2C) revealed distinct biological phases emerging during culture. Notably, transcripts associated with extracellular matrix (ECM) remodeling, such as MMP, are immediately upregulated after placing the explants into culture (Day 1) suggesting a rapid restructuring of the retinal microenvironment triggered by dissection and transfer to ex-vivo conditions. This agrees with our observation that mRNA level of some genes tends to alter faster than the others like those which are mainly related to injury. Early ECM remodeling is a well described component of neural tissue injury responses. It has been demonstrated that tissue handling, axotomy and altered diffusion gradients in retina can elicit a coordinated wound repair–like program involving ECM reorganization and gliosis ^9^,^12^,^14^. Interestingly we observed in our SLIM_GO analysis that the wound healing term has the highest GSVA score at Day 0. Importantly, a similar ECM- and gliotic remodeling has been described in glaucomatous optic neuropathy at the optic nerve head, where axonal stress is accompanied by changes in extracellular matrix organization and glial reactivity. This in parallel supports the interpretation that the Day 1 transcriptional signature reflects a biologically meaningful “injury-adaptation” response.

By Day 3, the transcriptional profile indicates broader cellular state transitions, including changes consistent with metabolic rewiring and attenuation of specialized neuronal/photoreceptor identity programs. For example, both our NGS and qPCR analysis demonstrates a downregulation of photoreceptor-associated genes such as OPN1SW (Figure 2C). Besides, GO terms such as nervous system process, photoreceptor cell maintenance, visual perception, and monoatomic ion transport are among the most downregulated ones in our GSVA analysis (Figure 3A). Importantly, we observed that neuronal/synaptic programs decrease early and remain suppressed during the culture according to our GO-Slim analysis time. Notably, we observed early enrichment of GO term “mitochondrial apoptotic pathway of intrinsic ligand” explaining early commence of death (starting from Day1) of RGC demonstrated by our TUNEL staining assay (Figure 2D). Consistent with our finding, there have been reports also suggesting that 50% of RGC were positively stained by TUNEL assay ^15^. Finally, a decline in mature neuronal and photoreceptor gene programs during organotypic maintenance has been reported previously in adult mouse retinal explants, where culture time has been associated with progressive remodeling of retinal morphology, reactive gliosis, and changes in neuronal circuitry and viability, even when lamination was preserved ^9, 12 16^.

Consistent with our findings, Tian et al. (2022) also reported downregulation of GO terms such as synapse organization, synaptic signaling, and ion channel activity, along with upregulation of apoptotic pathways, ribosomal biogenesis, and immune response in their FACS-sorted mouse RGCs in the range 1 to 3 days after injury in a mouse optic nerve crush model. Additionally, in agreement with our observation of upregulation of leadingedge genes including TLR2/4/7/8, MYD88, IL18, and IL1A in GSEA analysis, they also identified ATF3/CHOP transcription factors as the main driver of transcriptional changes in RGCs after injury and they also provide evidence that these TFs preferentially activate TLR signaling and neuroinflammation pathways. Thus, our data suggest that the same core injury-response transcriptional axis is engaged in porcine retinal tissue following enucleation, despite the differences in species and model system.

Ueno et al. (2018) reported GO terms such as upregulation of apoptotic process, immune system process, and downregulation of visual perception, and ion transport in their ONC model between day 1 and day 0 of gene expression analysis ^17^. They also found significant changes between day 4 and day 0, including upregulation of immune system and apoptotic processes, and downregulation of ion transport and visual perception. These findings are closely in agreement with our results from GSVA and GO-Slim analyses, highlighting the importance of these cellular processes during the early days of retina explant culture.

In Day 7 and Day 10, the appearance of cell-cycle regulators like E2F8 and CCNB3 suggests proliferation of non-neuronal populations such as microglia, Müller glia, astrocytes, and/or macrophages (Figure 2C). Retinal glia has been described previously to not only become reactive, but also proliferative under certain conditions of organotypic retinal cultures^11^,^12^, ^14^. This is particularly relevant as retinal explant cultures are frequently used as ex-vivo model system for injuries of the neuroretina. Comparative studies have demonstrated that key injury-related features such as RGC damage and glial activation are reproduced in retinal explant cultures, although the time kinetic and magnitude of responses may differ from those observed in in vivo axotomy models ^18^.

Importantly, dysregulation of retinal lipid and cholesterol homeostasis is increasingly linked to major retinopathies, including diabetic retinopathy (DR) and age-related macular degeneration (AMD), where altered cholesterol homeostasis and lipid-driven inflammation contribute to disease progression and linked to photoreceptor stress and degradation^19^,^20^,^21^,^22, 23^. The downregulation of the cholesterol biosynthetic process GO term in GSVA analysis consistent with a broader disruption of retinal cholesterol regulation. Additionally, Day 7 upregulation of OLR1 (LOX-1) is notable because LOX-1 is an oxidized-LDL scavenger receptor that can promote inflammatory endothelial responses, and it has been shown to mediate leukocyte recruitment in retinal vessels in an endotoxin-induced uveitis model.

Several lanes of our analysis indicate strong induction of prominent inflammatory state by Day 10. For instance, we observed induction of myeloid-associated genes (TLR8 and CLEC7A), strong enrichment of toll-like receptor signaling, chemotaxis, immune regulation, immune system process and inflammatory repones in our GSVA and GO-Slim analysis. Importantly, we observed substantial increase in the number of proliferating myeloid cells peaked at Day 7 and Day 10 (Figure 5A, B). Consistent with our findings, toll-like receptor activation and downstream cytokine induction are well documented in the context of retinal injury. Notably, in ischemia/reperfusion a rapid upregulation of TLR2/3, the adaptor MyD88 and inflammatory cytokines reflect the initiation of an innate immune danger response which is accompanied by microglial activation and is mechanistically in agreement with caspase-dependent neurodegeneration pathways ^24^. Importantly, in glaucoma, microglial activation and some cytokine/chemokine releases are increasingly recognized as modulators of RGC degeneration. Besides, the pronounced late-stage inflammatory signature observed in this study by GSVA and GSEA analysis is also conceptually consistent with the pathophysiology in glaucoma, where micro- and macroglia drive neuroinflammation are increasingly recognized as modulators of RGC vulnerability and disease progression even under controlled intraocular pressure. This parallel supports the notion that innate immune activation represents a convergent mechanism linking stress responses in ex-vivo retinal explants with those occurring in vivo during optic neuropathy ^25^,^26^,^27, 28^.

In parallel, multiple mitochondrial and oxidative phosphorylation related GO terms showed reduced activity at late time points (Figure 3A). Diminished mitochondrial electron transport and ATP synthesis terms are mechanistically plausible in ex-vivo retina, given the high metabolic demand of retinal neurons and the sensitivity of RGCs to bioenergetic perturbation. Our GSEA analysis provides orthogonal confirmation at the geneset level too. The late suppression of complex I assembly (Figure 4B), which we observed in our GSEA analysis, is especially intriguing in the context of neurodegeneration. Complex I deficits can reduce ATP availability and alter redox balance, promoting ROS and stress signaling in high-demand neurons. Mitochondrial dysfunction is strongly implicated across retinal degenerative conditions and optic neuropathies, including glaucoma, and is a focus of emerging therapeutic strategies ^25 29^. From a glaucoma/ONC perspective, this supports a model in which early innate immune activation and progressive bioenergetic compromise co-evolve during culture. These two processes are also intertwined in optic neuropathies, where glial inflammatory signaling and neuronal metabolic vulnerability jointly shape RGC fate ^30^.

Importantly, cell death/differentiation-related categories peak later during culture time in our GO-Slim analysis validating our GSVA analysis. These coordinated shifts recapitulate known features of retinal injury models, especially those involving axotomy. In vivo optic nerve crush (ONC) and optic nerve transection cause rapid RGC stress followed by progressive RGC loss, with prominent glial activation ^31^.

Porcine retinal explants are widely used as ex-vivo model systems of the retina where relevant aspects of the retinal structure are preserved for several days to weeks. However, due to the procedure of dissection - involving axotomy, disconnection from the systemic circulation and loss of RPE-photoreceptor connections - as well as culture conditions, initiate injury-driven and culture time-dependent cellular changes. As alterations in cell survival, inflammatory signaling, metabolic state and stress responses occur, the system must be understood as an ex-vivo neurodegeneration model in which changes unfold in a defined temporal sequence. Our time-resolved transcriptomic analysis demonstrates that acute injury and early adaptation—characterized by extracellular matrix (ECM) remodeling, stress responses, translation, and proteostasis—predominate between Day 1 and Day 3. This interval is optimal for investigating the molecular and cellular changes associated with neuronal retina injuries. In contrast, from Day 7 to Day 10, retinal explant cultures transition into a metabolically remodeled state distinguished by pronounced inflammation, rendering this period ideal for examining immune-related alterations and their impact in neuronal injury models. However, the early upregulation of cytokines in the injury model introduces additional complexity when employing this system for preventive studies. The biological context provided here will facilitate more accurate interpretation of experimental outcomes using the porcine retinal organotypic culture model.

## 4. Materials and Methods

### Porcine Retina Explant Culture

Enucleated porcine eyes were obtained from a local abattoir and transported on ice in sterile 0.9% NaCl, Globes were briefly disinfected by immersion in 70% (v/v) ethanol and transferred to a sterile Petri dish. Extraocular tissues were removed carefully without damaging the sclera. A circumferential incision was made just below the ciliary body (ora serrata), and the anterior segment (including lens/ciliary body/vitreous) was removed to yield the posterior eyecup. The eyecup was incised along the visual streak and unfolded to expose the neuroretina. Retinal explants were excised using a 4-mm biopsy punch, avoiding the visual streak. Punches were gently detached from the underlying RPE/choroid and transferred onto Millicell cell culture inserts pre-equilibrated (pore size 0.4 μm) with culture medium consisting of Neurobasal A medium, 2% B-27 supplement, 1% Glutamax, and 1% Antibiotic-Antimycotic (all from Gibco, Thermo Fisher Scientific). To avoid folding of the retina, a drop of media was placed on the membrane of the insert, and each punch was placed on a drop with the RGC facing the upper chamber of the insert. Two milliliters of media were added to the lower chamber of each well (6-well plate) to avoid floating of retina punches during the culture period. Explants were maintained at 37°C, 5% CO2, and 95% humidity, and the medium was replaced daily by transferring inserts into freshly prepared wells.

### RNA extraction and sequencing

For RNA extraction, retinal explants in 8 biological replicates per time point were generated as described above and transferred to 100 μL RLT buffer (RNeasy 96 Kit, Qiagen) supplemented with 1% β-mercaptoethanol and stored at −80 °C when required. Samples were homogenized using a pellet pestle motor (∼20 s) and processed using the RNeasy 96 plate-based protocol according to the manufacturer’s instructions. RNA quantity and integrity were assessed using a 96-channel Fragment Analyzer, applying a minimum RIN ≥ 7.5. For library preparation, 250 ng total RNA per sample (normalized to 25 ng/μL) was subjected to poly(A) enrichment (NEBNext Poly(A) Module) followed by stranded library construction (NEBNext Ultra II Directional). Libraries were quality-controlled and pooled equimolarly, then sequenced on an Illumina NovaSeq 6000 using paired-end 101 bp reads with dual indexing.

### Statistical analysis

Sample–sample similarity was quantified by Pearson correlation on variance-stabilized expression values (vstTransform) restricted to the top 5000 most variable genes across all samples. Correlations were computed using pairwise complete observations and visualized with ComplexHeatmap (legend range 0.8–1.0). Samples were unsupervised clustered using 1−r distance and complete linkage, with MFGroup annotation.

PCA was performed on the same VST matrix (top 5000 variable genes) using prcomp. Samples were colored by MFGroup, with 95% normal ellipses and labeled group centroids overlaid. The percentage of variance explained by PC1/PC2 was displayed on axes (45.78%, 19.7%).

Differential gene expression analysis was performed in DESeq2 using the raw count assay from exprMatrix (SummarizedExperiment). Samples were grouped by MFGroup (Day 0/1/3/7/10; Day 0 as reference) and analyzed with a paired design including EyeID (∼EyeID + MFGroup). Genes were retained if ≥10 counts in ≥3 samples. Wald tests were run with FDR (BH) α=0.05 and an effect-size threshold (lfcThreshold=1; |log2FC|≥1). Log2FC were optionally shrunk (apeglm). Pig gene symbols were mapped with biomaRt, and significant genes were summarized by baseline contrasts, visualized as Venn diagrams (ggVennDiagram), Up/Down count bars, and the top regulated genes per contrast.

Gene-set activity was estimated by GSVA on DESeq2 VST expression, using GO-BP gene sets with (minimum size ≥3, and Gaussian kernel). Time-point effects were tested on GSVA scores with limma R package (duplicateCorrelation and lmFit with EyeID as block), applying FDR ≤ 0.05. Significant GO terms were summarized as top-ranked contrasts and mapped to goslim_generic (BP; primary mapping) for heatmap visualization.

GSEA was performed with fgseaMultilevel on genes ranked by DESeq2 Wald statistic from a paired model (∼EyeID + MFGroup), after filtering (≥10 counts in ≥3 samples) and mapping Ensembl IDs to gene symbols. GO-BP gene sets (size 10–500) were tested per baseline contrast (Day 1/3/7/10 vs Day 0). Selected terms were visualized using plotEnrichment, the ranked-metric trace, and a leading-edge VST z-score heatmap (ComplexHeatmap).

### immunohistochemistry and immunofluoresence

Retinal tissues were fixed at 4 °C for up to 24 hours, processed using an automated tissue processor (Tissue-Tek VIP 6, Sakura), embedded in paraffin, and sectioned at 3 μm thickness. Hematoxylin and eosin (H&E) staining was performed with the Leica ST5020 Multistainer (Leica Biosystems, Germany) following standard protocols. Degenerating cells were detected by TUNEL assay (DIG-11-dUTP, Roche; TdT Enzyme, Promega) performed on the automated Leica Bond RX platform and visualized using the Bond Polymer Refine Detection Kit with 3,3′-diaminobenzidine (DAB) as the chromogen.

Immunofluorescence staining was performed on the automated Leica Bond platform using the Opal Multiplex IHC Kit (Akoya Biosciences). Heat-induced antigen retrieval and antibody stripping were carried out with Bond Epitope Retrieval Solutions 1 and 2 (ER1 and ER2, Leica Biosystems). Primary antibodies against Ki67, GFAP, and Iba1 were applied according to the staining sequence provided in (supplementary materials table1), followed by incubation with an HRP-labeled secondary antibody and Opal fluorescent reagent labeling. Nuclei were counterstained with spectral DAPI, and slides were mounted in ProLong Antifade Mounting Medium. Fluorescence imaging for analysis was conducted using an AxioScan 7 microscope (Zeiss), and quantitative analyses of Ki67+ and Iba1+ cells were performed using HALO Highplex Analysis software (Indica Labs). Representative pictures were imaged using a LSM900 (Zeiss).

### RNA extraction and qPCR

Tissue explants were generated in three biological replicates per time point and homogenized in RLT buffer with 1% ß-Mercaptoethanol as described above using a pellet pestle motor (∼20 s). Total RNA was isolated using the RNeasy Mini Kit (Qiagen) following the manufacturer’s instructions. RNA was eluted in RNase free water and stored at −80°C and RNA concentration was determined using a NanoDrop8000 (Thermo Fisher). For cDNA synthesis, RNA was reverse transcribed using the high-capacity cDNA Archive Kit (Applied Biosystems) according to the manufacturer’s instructions. Quantitative real time PCR (qPCR) was performed with two technical replicates using TaqMan chemistry using 20 ng cDNA and 2x TaqMan Fast Advanced MasterMix and 20x AoD Primer/Probe Mix. The list of TaqMan probes is provided in supplementary material Table2. Reactions were run on a QuantStudio 6 Flex system under fast cycling conditions (95°C 20-30 s, 40 cycles of 95°C 1 s, 60°C 20 s.). GAPDH was used as the housekeeping gene. Relative gene expression was determined using the 2^−ΔΔCT method. Fold change was calculated by dividing each gene’s expression level by the expression level of the corresponding Day 0 (control) sample. Subsequently, log^2^(fold change) values were computed for statistical analysis.

## Supplementary Materials

The following supporting information can be downloaded at: https://www.mdpi.com/article/doi/s1, Figure S1: adjacent_updown_counts_stacked_up.

## Author Contributions

S. K. wrote the manuscript and performed the data analysis and produced figures. R. B., G. G., and F. S. added input for publication strategy. F. S., T. S., P. G., and S. E. performed microscopy analysis. M.R., J. F., and performed qPCR experiment and analysis. S. B. and E. M have supervised analysis of RNA sequencing and data analysis. N. Z. orchestrated work packages.

## Funding

This work has been fully funded by Boehringer Ingelheim Pharma GmbH & Co.KG,.

## Data Availability Statement

Porcine RNA sequencing data generated in this study are deposited in GEO repository (accession number GSE318246).

## Acknowledgments

We thank Werner Rust, Nicole Ernst, Christoph Reuss and Alec Dick for their valuable support in porcine explant preparation, RNA preparation and sequencing. We also thank Martina Steinrock and Maria-Theresia Trinz for expert histology support.

During the preparation of this manuscript siavash khosravi used Copilot for the purposes of generating and proofreading text. Copilot has helped to debug the R code used for analysis. The authors have reviewed and edited the output and take full responsibility for the content of this publication.

## Conflicts of Interest

The authors declare no competing interests.

## Abbreviations

The following abbreviations are used in this manuscript:

AMD: age-related macular degeneration
B-27: B-27 supplement
BH: Benjamini–Hochberg (multiple-testing correction
BP: biological process
bp: base pairs
CO2: carbon dioxide
DESeq2: DESeq2 (R/Bioconductor package for RNA-seq differential expression
DGE: differential gene expression
DR: diabetic retinopathy
ECM: extracellular matrix
FDR: false discovery rate
GEO: Gene Expression Omnibus
GFAP: Glial fibrillary acidic protein
GO: Gene Ontology
GO-BP: Gene Ontology Biological Process
GO-: Gene Ontology Slim (high-level GO subset
Slim: 
GSEA: gene set enrichment analysis
GSVA: gene set variation analysis
Iba1: ionized calcium-binding adapter molecule 1
log2FC: log2 fold change
LOX-1: lectin-like oxidized low-density lipoprotein receptor 1
mRNA: messenger RNA
NaCl: sodium chloride
NES: normalized enrichment score
NF-κB: nuclear factor kappa B
ONC: optic nerve crush
PCA: principal component analysis
RGC: retinal ganglion cell
RIN: RNA integrity number
RLT: RLT lysis buffer (Qiagen
RNA: ribonucleic acid
RNA-: RNA sequencing
seq: 
RPE: retinal pigment epithelium
VST: variance-stabilizing transformation

## Disclaimer/Publisher’s Note

The statements, opinions and data contained in all publications are solely those of the individual author(s) and contributor(s) and not of MDPI and/or the editor(s). MDPI and/or the editor(s) disclaim responsibility for any injury to people or property resulting from any ideas, methods, instructions or products referred to in the content.

## Notes

### Competing Interest Statement

The authors have declared no competing interest.

