## Supplementa file for "Transcriptomic analysis of organotypic porcine retina cultures"

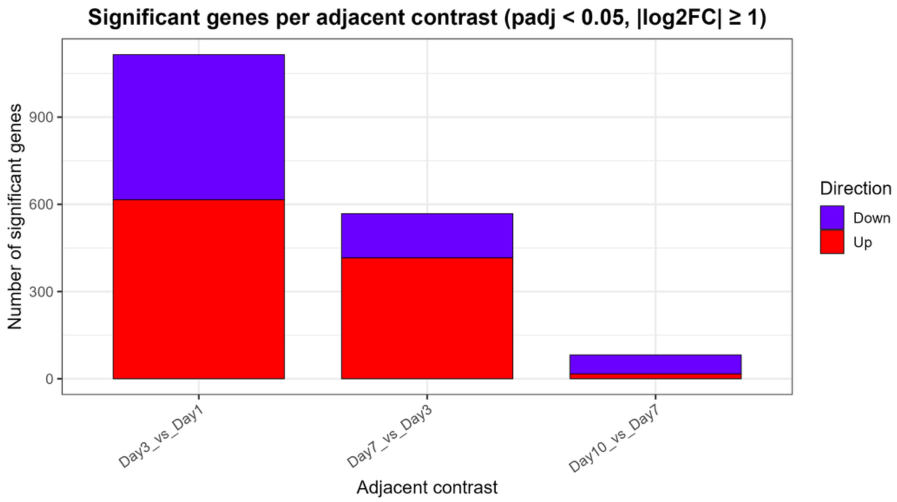


*Supplementary Figure 1:The number of significant genes in adjacent contrasts decreases during the culture period. Related to Figure 2A.*

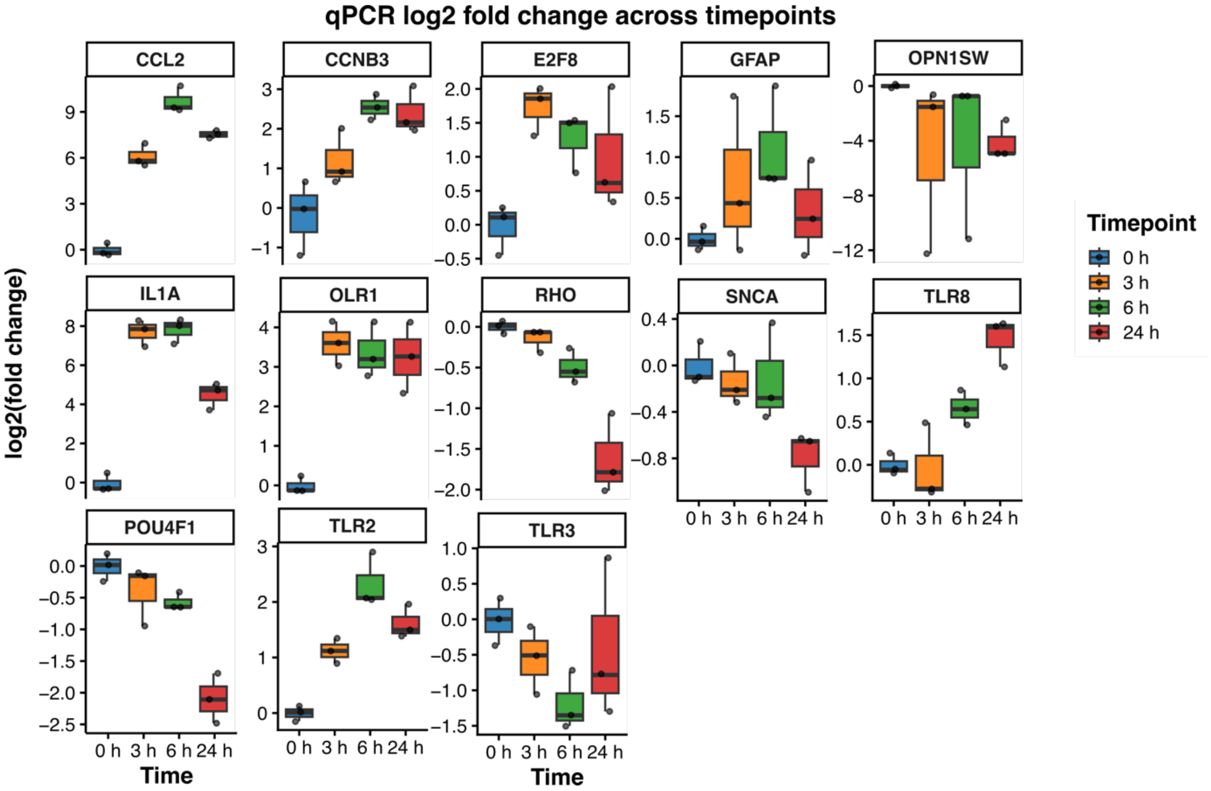


*Supplementary Figure 2: mRNA levels of genes alter differentially in the first 24 hours after dissections. Related to Figure 2.*

mRNA levels of selected genes were measured at 1, 3, 6, and 24 hours post-dissection.

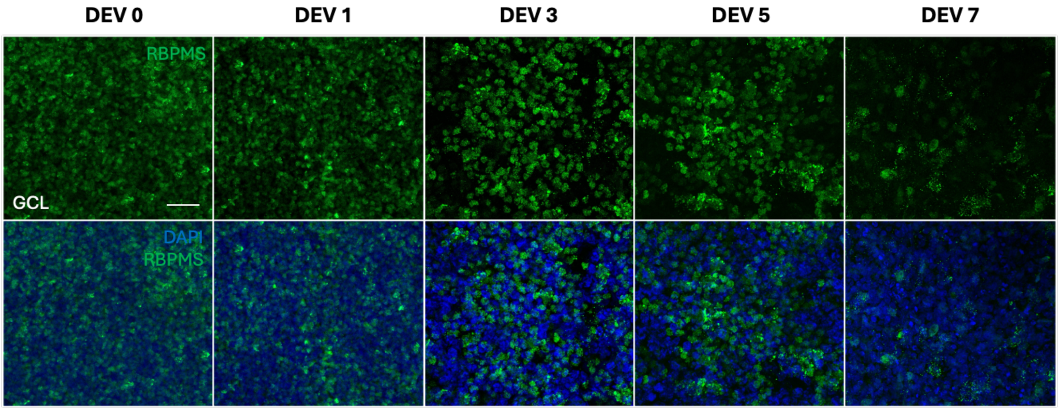


*Supplementary Figure 3 Number of retinal ganglion cells reduces during the culture time. Related to Figure 5.*

Porcine retina flat mounts were fixed, stained with anti-RBPMS antibody and DAPI, and analyzed via microscopy. The top panel displays RBPMS staining; the lower panel shows both DAPI and RBPMS staining (DEV = Day Ex Vivo).

*Supplementary Table 1: Antibodies used in this study listed in staining order*

| Marker | Cat#, Vendor | Concentration or dilution | Antigen retrieval / antibody stripping | Fluorescent reagent |
|---|---|---|---|---|
| Ki67 | ab16667, abcam | 0.31 µg/ml | ER1 (pH6, 95°C/ 30 min) | OPAL 690 |
| GFAP | ab7260, Abcam | 1:8000 | ER2 (pH9, 95°C/ 20 min) | OPAL 570 |
| Iba1 | 019-19741, Wako | 0.5 µg/ml | ER2 (pH9, 95°C/ 20 min) | OPAL 520 |
| RBPMS | NBP2-20112-NovusBio | 1 to250 | none | |

*Supplementary Table 2: Gene probes (Thermo Fisher Scientific) for qPCR*

| Gene Symbol | Assay ID |
|---|---|
| GAPDH | Ss03375629_u1 |
| GFAP | Ss03373547_m1 |
| CCL2 / MCP1 | Ss03394377_m1 |
| POU4F1 | Ss02916785_m1 |
| OPN1SW | Ss03393999_m1 |
| SNCA | Ss03379644_u1 |
| RHO | Ss03394397_m1 |
| IL1A | Ss03391335_m1 |
| E2F8 | Ss06897499_s1 |

| | |
|---|---|
| CCNB3 | Ss06916241_m1 |
| TLR8 | Ss03383235_u1 |
| TLR2 | Ss03381278_u1 |
| TLR3 | Ss03388861_m1 |
| OLR1 | Ss03392357_m1 |

### Material and methods:

Porcine retinal explant preparation and immunohistochemistry:

Porcine eyes were surface-sterilized by immersion in 70% ethanol and dissected under sterile conditions. The anterior segment and vitreous were removed, and the posterior eye cup was flattened by four radial incisions. Retinal punches of 4 mm in diameter were obtained using a biopsy corer, avoiding the visual streak region to ensure tissue homogeneity. Individual explants were placed onto semipermeable membrane inserts in 6-well plates (ganglion cell layer facing upward) and cultured in Neurobasal-A medium supplemented with B27 (2%), GlutaMAX (1%), and Antibiotic-Antimycotic (1%) at 37°C and 5% $CO_2$. At designated time points day 0, 1, 3, 5, and 7, explants were fixed with 4% paraformaldehyde for 1 h at room temperature and processed as whole-mount preparations. For immunofluorescence, explants were permeabilized and blocked in 2% BSA / 2% Triton X-100 in PBS for 1 h at room temperature, then incubated with a rabbit anti-RBPMS primary antibody (NovusBio, NBP2-20112; 1:250) in blocking buffer for 48 h at 4°C. After three washes in PBS, explants were incubated overnight at room temperature in the dark with a donkey anti-rabbit Alexa Fluor 488 secondary antibody (Molecular Probes, A21206; 1:1000) together with DAPI (1:1000). Following additional PBS washes, flatmounts were mounted with Aqua Poly/Mount and imaged.
